# PD-1 blockade exacerbates *Mycobacterium tuberculosis* infection in rhesus macaques

**DOI:** 10.1101/2020.08.05.237883

**Authors:** Keith D Kauffman, Shunsuke Sakai, Nickiana E Lora, Sivaranjani Namasivayam, Paul J Baker, Olena Kamenyeva, Taylor W Foreman, Christine E Nelson, Deivide Oliveira-de-Souza, Caian L. Vinhaes, Ziv Yaniv, Cecilia S Lindestam Arleham, Alessandro Sette, Gordon J Freeman, Rashida Moore, the NIAID/DIR Tuberculosis Imaging Program, Alan Sher, Katrin D Mayer-Barber, Bruno B Andrade, Juraj Kabat, Laura E Via, Daniel L Barber

**Affiliations:** T Lymphocyte Biology Section, Laboratory of Parasitic Diseases, National Institute of Allergy and Infectious Diseases, National Institutes of Health, Bethesda, Maryland, USA; Immunobiology Section, Laboratory of Parasitic Diseases, National Institute of Allergy and Infectious Diseases, National Institutes of Health, Bethesda, Maryland, USA; Inflammation and Innate Immunity Unit, Laboratory of Clinical Immunology and Microbiology, National Institute of Allergy and Infectious Diseases, National Institutes of Health, Bethesda, USA; Biological Imaging Section, Research Technologies Branch, National Institute of Allergy and Infectious Diseases, National Institutes of Health, Bethesda, Maryland, USA; Multinational Organization Network Sponsoring Translational and Epidemiological Research (MONSTER) Initiative, Intituto Gonçalo Moniz, Fundação Oswaldo Cruz, Salvador 40296-710, Brazil; Office of Cyber Infrastructure and Computational Biology, National Institute of Allergy and Infectious Diseases, National Institutes of Health, Bethesda, Maryland, USA; Division of Vaccine Discovery, La Jolla Institute for Immunology, La Jolla, California, USA; Department of Medicine, University of California San Diego, La Jolla, California, USA; Department of Medical Oncology, Dana-Farber Cancer Institute, Harvard Medical School, Boston, MA 02115, USA; Comparative Medicine Branch, National Institute of Allergy and Infectious Diseases, National Institutes of Health, Bethesda, Maryland, USA; Division of Intramural Research, National Institute of Allergy and Infectious Diseases, National Institutes of Health, Bethesda, Maryland, USA; Tuberculosis Research Section, Laboratory of Clinical Infectious Diseases, National Institute of Allergy and Infectious Diseases, National Institutes of Health, Bethesda, Maryland, USA

## Abstract

Boosting immune cell function by targeting the co-inhibitory receptor PD-1 may have applications in the treatment of chronic infections. Here we examine the role of PD-1 during *Mycobacterium tuberculosis* (Mtb) infection of rhesus macaques. Animals treated with αPD-1 mAb developed worse disease and higher granuloma bacterial loads compared to isotype control treated monkeys. PD-1 blockade increased the number and functionality of granuloma Mtb-specific CD8 T cells. In contrast, Mtb-specific CD4 T cells in αPD-1 treated macaques were not increased in number or function in granulomas, upregulated high levels of CTLA-4 and exhibited reduced intralesional trafficking in live imaging studies. In granulomas of αPD-1 treated animals, multiple pro-inflammatory cytokines were elevated, and more cytokines correlated with bacterial loads, leading to the identification of a role for caspase 1 in the exacerbation of tuberculosis after PD-1 blockade. Lastly, increased Mtb bacterial loads after PD-1 blockade were found to associate with the composition of the intestinal microbiota prior to infection in individual macaques. Therefore, PD-1-mediated co-inhibition is required for control of Mtb infection in macaques, perhaps due to its role in dampening detrimental inflammation as well as allowing for normal CD4 T cell responses.

## INTRODUCTION

*Mycobacterium tuberculosis* (Mtb) infection is the leading cause of death due to a single infectious agent worldwide, despite the availability of antibiotics that can effectively treat most Mtb infections (*1*). Drugs that target the host rather than the bacteria, i.e. host-directed therapies (HDTs), may be useful in shortening the standard 6-month long course of antibiotic treatment, as well as provide sorely needed new options for the treatment of drug resistant infections (*2–4*). In particular, there is interest in developing strategies to boost host-protective immune responses, or on the other hand limit the detrimental inflammation that causes tissue destruction and promotes bacterial growth during tuberculosis. However, the mechanisms of host resistance and tissue pathology during Mtb infection are incompletely understood, impeding the development of HDTs.

PD-1 is a co-inhibitory receptor primarily expressed on activated CD4 and CD8 T cells that has been shown to limit the function of pathogen-specific T cells during chronic infection and tumor-specific T cells during cancer (*5, 6*). Importantly, blockade of the PD-1 receptor or its ligands with monoclonal antibodies enhances the number and function of anti-tumor cytotoxic T cells resulting in greatly enhanced tumor control, and there are currently multiple PD-1 targeting drugs approved for use against various malignancies (*6*). The major success of such immune checkpoint blockade targeting drugs in cancer treatment has highlighted how potent such approaches can be in the treatment of human disease. Indeed, boosting T cell function by blocking PD-1 is often suggested as a therapy for tuberculosis (TB) (*7*).

Human Mtb-specific T cells in circulation can express low levels of PD-1 during disease, and in vitro blockade of PD-1 can enhance T cell responses, although the effects are modest (*8*). The first in vivo data on the role of PD-1 in Mtb infection came from knock out mouse studies where it was found that PD-1^-/-^ mice die very rapidly after Mtb infection compared to WT mice (*9, 10*). In the absence of PD-1, CD4 T cells and, to a lesser extent, CD8 T cells drive this early mortality (*10*). Although the T cell mechanisms that cause pathology in Mtb infected PD-1^-/-^ mice are not completely understood, we have shown that the over production of IFNγ by CD4 T cells is at least partly responsible (*11*), and in a human in vitro 3D-granuloma model, it was found that PD-1 blockade drives higher bacterial loads in a TNF-dependent manner (*12*). Consistent with these data showing a host-protective role for PD-1 in Mtb infection, clinical case reports of checkpoint blockade-associated tuberculosis in patients treated with anti-PD-1 are accumulating in the literature (*12–18*). *Mycobacterium avium* infections have also been observed in individuals undergoing PD-1 blockade cancer immunotherapy (*19*). This has led to the hypothesis that the negative regulation of T cells through PD-1 is required for optimal control of Mtb infection, perhaps by the inhibition of detrimental hyperinflammatory T cell responses. However, due to the caveats inherent to the murine model of TB and the anecdotal nature of the published case reports, the role of PD-1 in host-resistance to Mtb infection in vivo has yet to be definitively addressed.

Macaques are considered the gold standard pre-clinical model of Mtb infection, as they very accurately recapitulate many features of human tuberculosis pathogenesis and immune responses against the pathogen (*20*). Therefore, in the current study we treated Mtb infected rhesus macaques with αPD-1 to examine the role of this clinically important regulatory pathway during tuberculosis. We find that PD-1 blockade drove higher levels of inflammation and increased bacterial loads in pulmonary granulomas. Taken together with the previously published data, these results strongly indicate that, somewhat counter-intuitively, the co-inhibitory receptor PD-1 is host protective during Mtb infection and that negative immune regulation can be an important aspect of host resistance during tuberculosis.

## RESULTS

### Increased bacterial loads after PD-1 blockade

PD-1 expression was most highly expressed on antigen (Ag)-specific CD4 and CD8 T cells in lung granulomas compared to other tissues, including the blood and even the bronchoalveolar lavage (BAL) (Figure 1a, b). In granulomas, PD-1 staining was largely restricted to the peripheral lymphocyte-rich cuff (Figure 1c) and was detectable on CD4+ as well as CD8+ cells (Figure 1d). Therefore, PD-1 is preferentially expressed on Mtb-specific T cells in pulmonary granulomas, the primary site of bacterial replication. To examine the outcome of PD-1 blockade, macaques were treated with primatized anti-PD-1 or rhesus macaque isotype control starting two weeks post-infection (Figure 2a). Our hypothesis was that PD-1 blockade would impair control of the infection. Accordingly, a dose and strain of Mtb was chosen that, in our experience, results in a slowly progressive course of infection, so that the virulence of the bacteria would not mask the predicted effect of PD-1 blockade.

**Figure 1.**
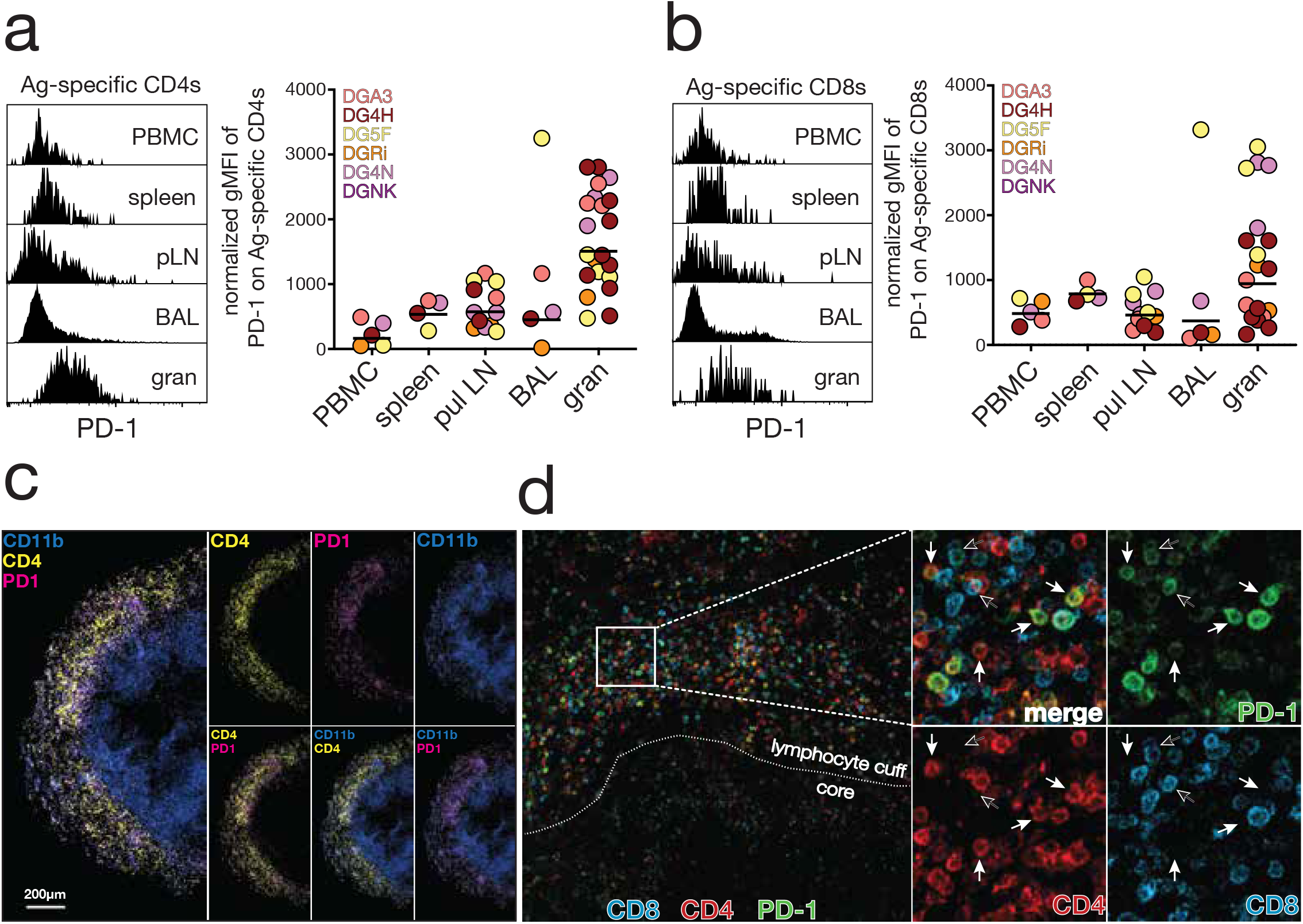
PD-1 expression is highest on T cells in pulmonary granulomas. a-b) Example histograms and summary graphs of PD-1 staining by flow cytometry on (a) CD4 T cells and (b) CD8 T cells from the indicated tissues at necropsy, ~15 weeks post infection with 30-50 CFU of H37Rv-mCherry. Mtb-specific T cells identified as TNF+ by intracellular cytokine staining after stimulation with a combination of CD4 and CD8 T cell Mtb peptide megapools. The normalized geoMFI was calculated by subtracting the geoMFI of PD-1 negative T cells in blood of the same animal from the TNF+ Ag-specific T cells in different tissues. c) PD-1 (magenta), CD4 (yellow), and CD11b (blue) staining in an Mtb granuloma. Left image is merged and smaller images on right show individual staining (top) and two-stain combinations (bottom). Image is from animal DG4H. Note that the mCherry signal was undetectable at this time point. d) Costaining of PD-1 (green), CD4 (red), and CD8 (blue) in a granuloma. Left image is merged and shows the boundary of the lymphocyte cuff and core. Expanded image shows individual stains and merged image. White arrows indicate PD-1^+^ CD4^+^ cells and black arrows indicate PD-1^+^CD8^+^ cells. This image from animal MFN is from a separate infection (five weeks post-infection with 12-14 CFU Mtb-Erdman) and is included as CD8 staining in granulomas was not performed in PD-1 blockade study.

**Figure 2.**
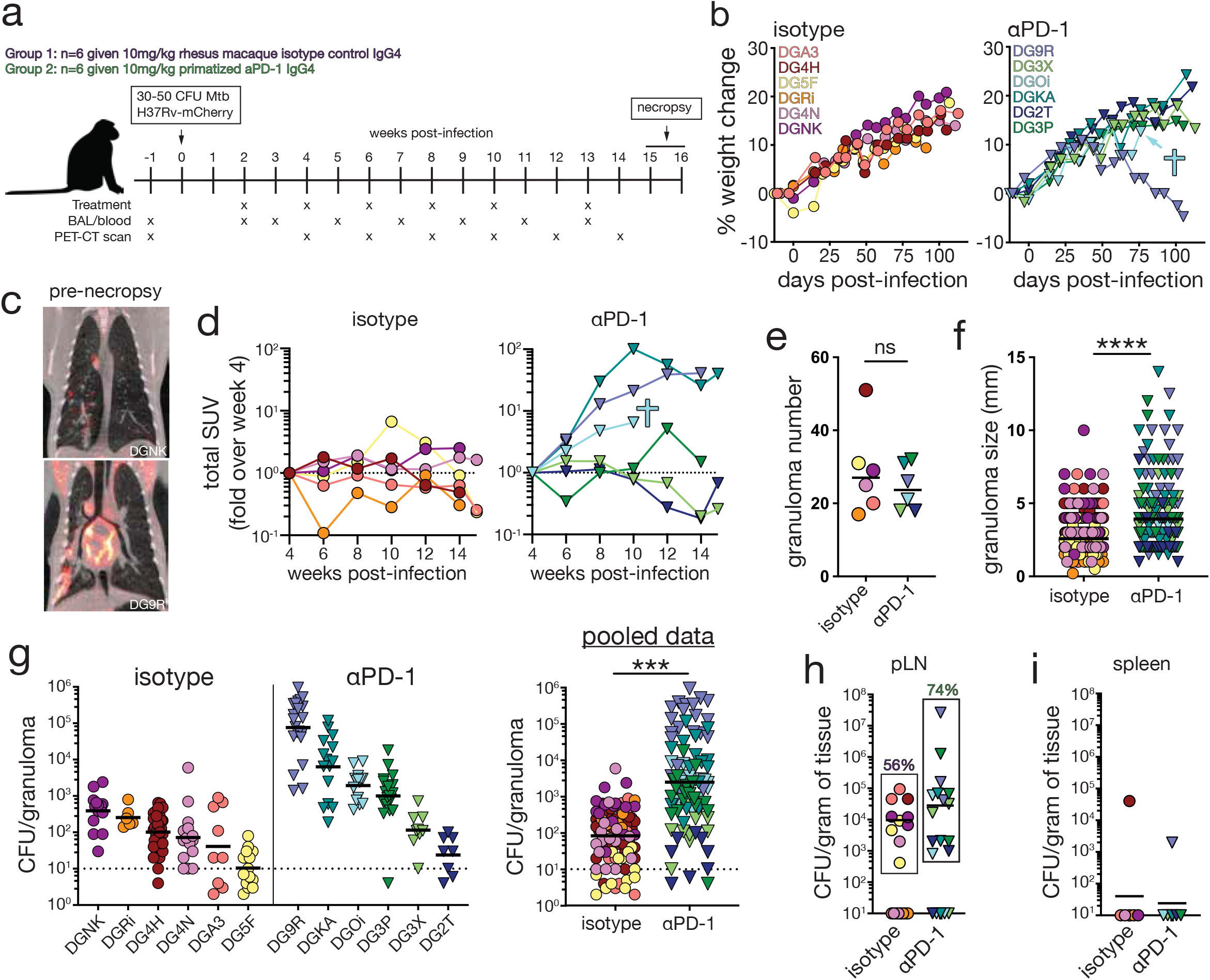
PD-1 blockade exacerbates tuberculosis in macaques. a) Twelve animals were infected with 30-50 colony forming units (cfu) of H37Rv-mCherry. Starting two weeks post-infection, animals were treated with a rhesus macaque IgG4 isotype control (n=6) or a primatized αPD-1 mAb (n=6). Animals were necropsied at weeks 15 or 16 post-infection. b) Weight change of animals receiving isotype control (left) and anti-PD-1 (right). Blue cross indicates euthanasia of DGOi prior to study due to the development of severe pulmonary distress. c) Example PET-CT image from isotype control (top) or αPD-1 (bottom) treated animals. d) Fold change over week 4 value of total lung standardized uptake value (SUV) in isotype (left) and αPD-1 treated (right) animals. e) At necropsy the entire lung was closely examined, and all granulomas were isolated. Graph represents the number of granulomas recovered from the whole lung. f) Dimeter of each granuloma was measured at necropsy. Average of two perpendicular measurements was taken for irregularly shaped lesions. g) Bacterial loads in granulomas displayed in individual animals (left) or pooled by experimental group (right). Each dot represents an individual granuloma. Dotted line indicates limit of detection. h-i) Bacterial burden in (h) pulmonary lymph nodes and (i) spleen. Percentage of total lymph nodes from each experimental group that had positive bacterial cultures is indicated.

All six isotype control treated animals survived until the pre-determined endpoint (week 15-16) and displayed no signs of weight loss and had stable disease as measured by ^18^FDG-PET-CT imaging (Figure 2b-d). In contrast, one of the αPD-1 treated animals (DGOi) developed acute symptoms and was euthanized early, another developed cachexia and relatively elevated ^18^FDG uptake but survived until the study end (DG9R), and yet another displayed higher PET-CT scores but did not lose weight (DGKA) (Figure 2b-d). Upon necropsy, we found that isotype control and αPD-1 treated animals had similar numbers of granulomas, but the granulomas in αPD-1 treated macaques were larger (Figure 2e-f). Importantly, αPD-1 treated macaques had an ~20-fold increase in the numbers of bacteria in their granulomas relative to animals treated with isotype control mAb (Figure 2g). While 4 of the 6 αPD-1 treated macaques displayed increased bacterial loads, two animals (DG3X and DG2T) were “non-responders” and had low bacterial loads similar to the isotype control treated animals. Although the bacterial loads were similar in the lung draining lymph nodes, a higher percentage of pulmonary lymph nodes were infected in the αPD-1 treated group compared to control macaques, and most spleen samples did not contain bacteria (Figure 2h-i). Thus, PD-1 blockade increased bacterial loads in pulmonary granulomas but did not result in disseminated infection.

### Frequency of Mtb-specific T cells in granulomas

The magnitude of the Mtb-specific CD4 and CD8 T cell responses were enumerated by intracellular staining after stimulation with both CD4 and CD8 Mtb peptide megapools (*21, 22*). Rather than the increased T cell expansions expected to result from PD-1 blockade, we found that the accumulation of Mtb-specific CD4 T cells in the blood and airways was delayed by PD-1 blockade, while the kinetics of Mtb-specific CD8 T cell responses in the BAL fluid were similar to control animals (Figure 3a-b). The frequency of Mtb-specific CD4 and CD8 T cells in the pulmonary lymph nodes at necropsy were not affected by PD-1 blockade (Figure 3c, f). In granulomas, the frequency of Mtb-specific CD4 T cells was similar between isotype control and αPD-1 treated animals, while Mtb-specific CD8 T cells were significantly increased by PD-1 blockade (Figure 3d, g). These results are in contrast to studies of Mtb infection in PD-1 KO mice, where it was shown that Mtb-specific CD4 T cells in the lungs are significantly increased by PD-1 deficiency and Mtb-specific CD8 T cells were much less effected (*10*).

**Figure 3.**
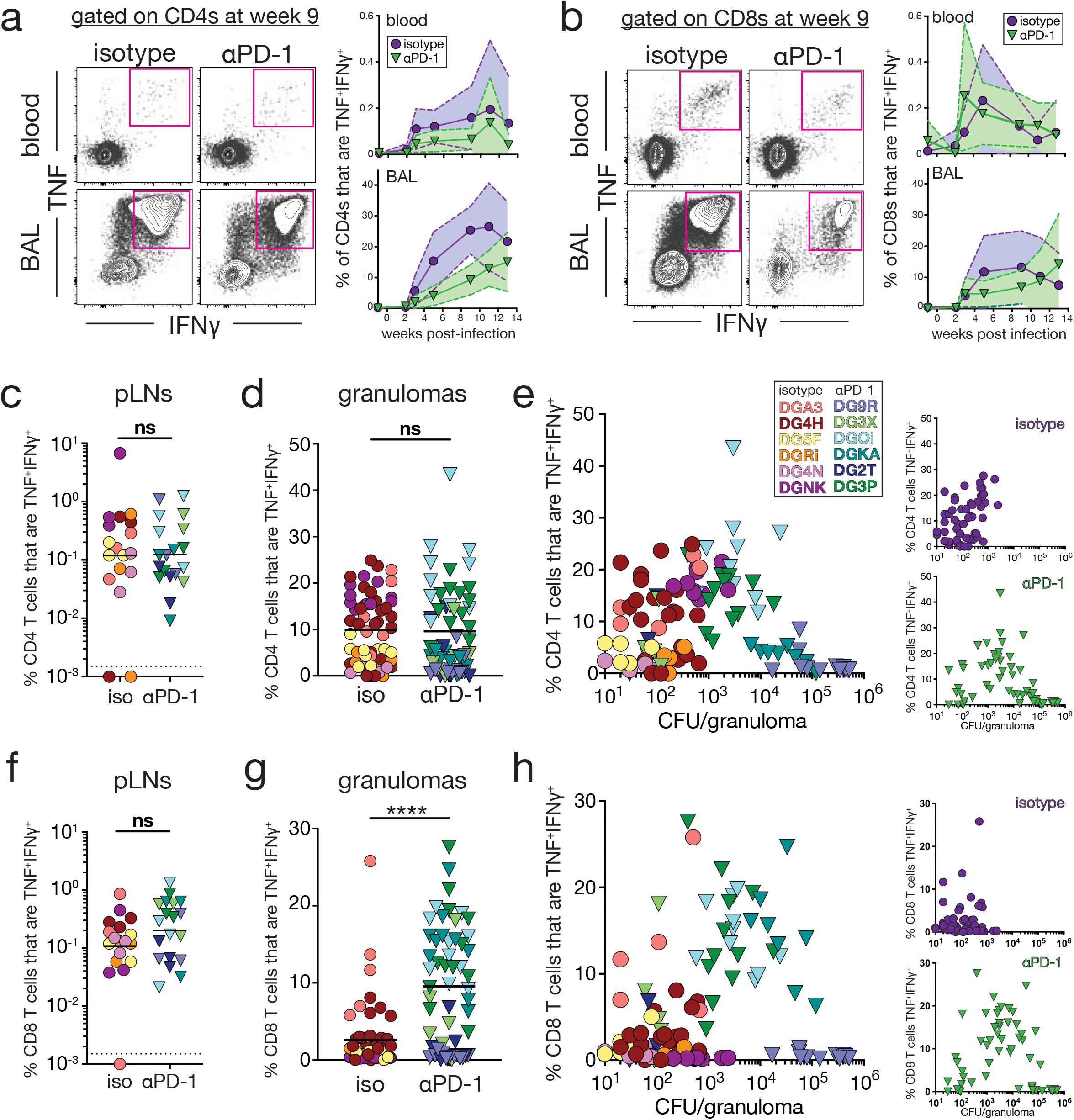
PD-1 blockade increases Mtb-specific CD8 T cell responses in granulomas. a) Example FACS plots and summary graphs of CD4 T cells from the blood and BAL following in vitro stimulation with the MTB300 peptide megapool. b) Example FACS plots and summary graphs of CD8 T cells from the blood and BAL following in vitro stimulation with an Mtb MHC-I peptide megapool. c-d) Graphs showing the percentage of CD4 T cells in the pulmonary lymph nodes (c) and granulomas (d) that are TNF+IFNγ+ following in vitro stimulation. e) Comparison of the bacterial burden of each granuloma with the percent of CD4 T cells that are TNF+IFNγ+ following in vitro stimulation. Small insets display the same information separated by experimental group with isotype control (top) and αPD-1 (bottom). f-g) Graphs showing the percentage of CD8 T cells in the pulmonary lymph nodes (f) and granulomas (g) that are TNF+IFNγ+ following in vitro stimulation. h) Comparison of the bacterial burden of each granuloma with the percent of CD8 T cells that are TNF+IFNγ+ following in vitro stimulation. Small insets display the same information separated by experimental group with isotype control (top) and αPD-1 (bottom).

Interestingly, PD-1 blockade altered the relationship between the magnitude of the Mtb-specific T cell response and bacterial loads. At bacterial loads lower than ~2000-5000 CFU/granuloma, there was trend for a positive correlation between Ag-specific T cells and numbers of bacteria, but at higher bacterial loads this switched to a negative correlation (Figure 3e, h). In other words, Ag-specific T cell responses were the highest in granulomas near the middle of the CFU distribution, and the granulomas with the lowest CD4 T cell responses were at either the lowest or highest end of the CFU distribution. We previously observed a nearly identical non-monotonic relationship between Ag-specific CD4 T cells and bacterial loads in an independent group of macaques that developed a similarly wide range of granuloma bacterial loads after infection with a more virulent strain of Mtb (*23*). Therefore, we suggest that this complex relationship between bacterial loads and T cell responses in individual granulomas is not related to PD-1 blockade itself but may be a previously unrecognized normal feature of the heterogeneity in tuberculosis granulomas.

PD-1 blockade had no effect on the frequency of Foxp3+ CD4 T cells, MAIT cells or γδT cells in the blood and BAL at any time point after infection, or in granulomas at necropsy (Supplemental Figure 1). Overall these results show that PD-1 blockade results in greater Mtb-specific CD8 but not CD4 T cell responses in pulmonary lesions at the time of necropsy. We should point out, however, that we do not know how PD-1 blockade may have impacted T cell responses in granulomas at earlier timepoints.

### Function of Mtb-specific T cells in granulomas

We next examined the effect of PD-1 blockade on T cell functionality. Mtb-specific CD4 T cells were identified as TNF+ after stimulation with peptide megapools. PD-1 blockade did not change the production of IFNγ, CD153 or granzyme B by Mtb-specific CD4 T cells but resulted in a slight reduction in IL-2 and increased IL-17A production (Figure 4a). Overall there was very little difference in the functional profile of Mtb-specific CD4 T cells. However, we observed a striking increase in the expression of CTLA-4 on TNF+ Mtb-specific CD4 T cells in the granulomas of animals treated with αPD-1 compared to the isotype control mAb (Figure 4b). This was also observed in infected PD-1 KO mice, where CTLA-4 is upregulated on Mtb-specific CD4 T cells in the lungs (*10*). In contrast to CD4 T cells, TNF+ Mtb-specific CD8 T cells in granulomas displayed a significant increase in production of IFNγ, IL-2 and granzyme B and did not have elevated CTLA-4 expression (Figure 4c-d). Therefore, PD-1 blockade did not increase the number or the functional potential of Mtb-specific CD4 T cells, and instead resulted in increased expression of another negative regulatory molecule. In contrast, PD-1 blockade resulted in increases in the number and function of Mtb-specific CD8 T cells. It is important to point out, however, that this analysis measures the potential of the cells to respond to high level peptide restimulation in vitro, and actual cytokine production may be increased in vivo by PD-1 blockade without major changes to their maximum secretion capacity upon high level peptide stimulation ex vivo. For this reason, we therefore next measured soluble mediators in granuloma homogenates.

**Figure 4.**
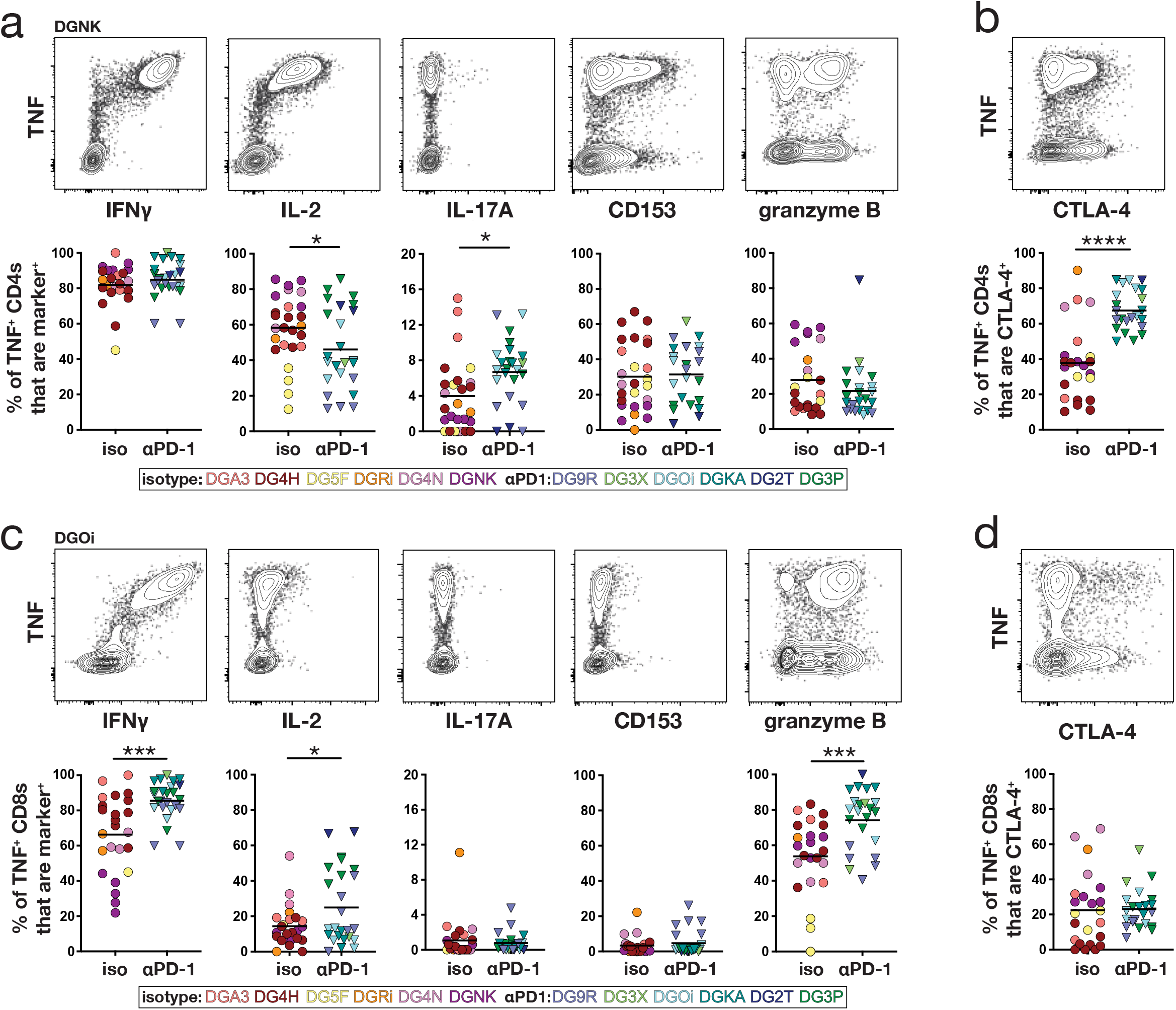
Enhanced function of Mtb-specific CD8 T cells after PD-1 blockade. a) Example FACS plots of cytokine staining on total CD4 T cells of a granuloma from animal DGNK and summary graphs of individual cytokines from TNF+ CD4 T cells. Each symbol is a granuloma from indicated animal. b) Example FACS plot of CTLA-4 and TNF staining on total CD4 T cells of a granuloma from animal DGNK and summary graph of percent CTLA-4+ of TNF+ CD4 T cells. c) Example FACS plots of cytokine staining on total CD8 T cells of a granuloma from animal DGOi and summary graphs of individual cytokines from TNF+ CD8 T cells. Each symbol is a granuloma from indicated animal. d) Example FACS plot of CTLA-4 and TNF staining on total CD4 T cells of a granuloma from animal DGNK and summary graph of percent CTLA-4+ of TNF+ CD4 T cells.

### Increased inflammation in granulomas after PD-1 blockade

We first profiled soluble mediators in plasma and granuloma homogenates. Interestingly, starting ~1 week after the first antibody infusion, there was a large burst of soluble CD40L and CCL3 in plasma of all six αPD-1 treated animals, but these molecules were undetectable in all of the isotype control treated animals (Supplemental Figure 2). Moreover, there was a strong correlation between soluble CD40L and CCL3 in the plasma (Supplemental Figure 2), which may be indicative of systemic T cell activation after PD-1 blockade. We did not perform blockade in uninfected macaques, so it is not clear if PD-1 blockade would have resulted in systemic CD40L and CCL3 detection in the absence of Mtb infection.

In granulomas, the three cytokines that were the most upregulated after αPD-1 treatment were IL-18, IFNγ, and TNF (Figure 5a). This is consistent with mouse model data indicating that IFNγ promotes pathology in Mtb infected PD-1^-/-^ mice and with in vitro data showing that TNF can promote increased bacterial growth after PD-1 blockade in an in vitro human 3D granuloma model (*11, 12*). Sparse canonical correlation analysis showed that the quality of inflammation was greatly impacted by PD-1 blockade, with IL-18 a major driver of differences between the two groups (Figure 5b). Further analysis of the Spearman correlations in the cytokine network found that IFNγ was the largest node of the cytokine network in granulomas isolated from control animals, whereas IL-18 was the most interconnected node in αPD-1 treated animals (Figure 5c). Overall, the density of cytokine network was much higher in granulomas from αPD-1 compared to isotype control treated animals. The increased cytokine concentrations and interconnectivity of the cytokine networks collectively indicate that the granulomas from the αPD-1 treated animals are more inflamed.

**Figure 5.**
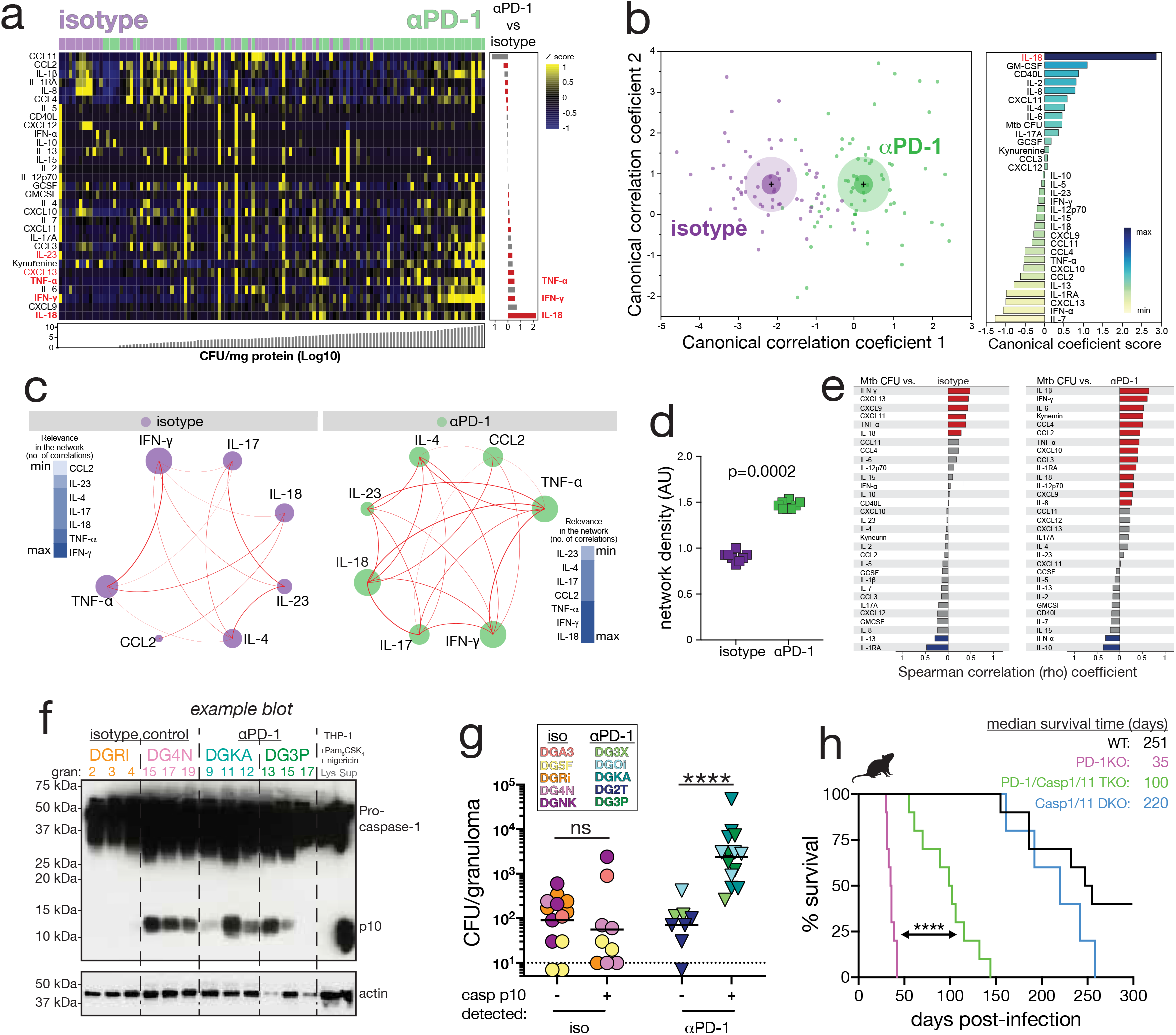
Increased inflammation in granulomas after PD-1 blockade. Soluble mediators were measured in granuloma homogenate supernatants of isotype control and αPD-1 treated macaques. Granuloma homogenate samples from DG9R and DG4H were not available and are not included in the cytokine multiplex analysis. a) *Left panel:* Data were log-transformed and z-score normalized. Markers were ranked according to mycobacterial load values (CFU/mg of protein) and fold-difference value between the different experimental groups. A heatmap was built to describe the overall expression profile of the inflammatory markers in each granuloma from monkeys labeled according to the experimental group indicated by distinct colors (green for isotype control and purple for anti-PD1). *Right panel:* Average fold-difference values in expression of inflammatory markers in granulomas from monkeys treated with anti-PD1 vs. isotype controls are described (log-transformed values). Differences which reached statistical significance with the Mann-Whitney *U* test adjusted for multiple comparisons using the Holm-Bonferroni’s method (Adjusted P < 0.05) are represented in colored bars. b) *Left panel:* Discrimination of groups using combination of inflammatory biomarkers per granuloma. In an exploratory approach, a sparse canonical correlation analysis (sCCA) was employed to test whether experimental groups could be distinguished based on the overall expression profile of all the markers measured as well as mycobacterial loads (*M. tuberculosis* CFU values per mg of protein). *Right panel:* Canonical coefficient scores were calculated to identify the biomarkers most responsible for the difference between groups in the sCCAmodel. c) Spearman correlation matrices of the biomarker expression levels in each study group were built and Circos plots were used to illustrate the correlation networks. Each circle represents a different plasma parameter. The size of each circle is proportional to the number of significant correlations. The connecting lines represent statistically significant correlations (p<0.01). Red connecting lines represent positive correlations while blue lines infer negative correlations (no negative correlations were found). Color intensity is proportional to the strength of correlation (rho value). Node analysis was used to illustrate the number of significant correlations per marker. Markers were grouped according to the number of connections from minimum to maximum numbers detected. d) Network density was compared between the groups using permutation test (100 permutations). e) Spearman correlation analysis was used to test association between cytokine values in granulomas and the mycobacterial loads (CFU/mg of protein) in monkeys receiving isotype controls (left panel) and in those who were treated with anti-PD-1 (right panel). Bars represent the Spearman rank (rho) values. Colored bars indicate statistically significant correlations (p < 0.05) after adjustment for multiple measurements. Red color infers positive correlation whereas blue color denotes negative correlations and grey bars are nonsignificant associations. f) Example Western blot of caspase-1 in granuloma homogenates. See Supplemental Figure 2 for all blots ran for this analysis. g) Bacterial loads in granulomas with detectable active caspase-1 p10 or not in isotype control or αPD-1 treated macaques. h) Survival curves and MST of WT, PD-1KO, caspase 1/11 double KO, or PD-1/caspase1/11 triple KO mice after aerosol Mtb-H37Rv infection. Data are pooled from two independent survival studies.

We next looked for correlations between different soluble mediators and bacterial loads. There were more positive correlations between soluble mediators and bacterial loads in granulomas of αPD-1 treated animals (14 of 31 analytes) compared to isotype control treated animals (6 of 31 analytes) (Figure 5e). Increased levels of IL-18, IFNγ, and TNF positively correlated with bacterial loads in both experimental groups. Several molecules correlated with CFU only in αPD-1 but not isotype control treated animals, including IL-1β, IL-1RA, IL-6, IL-8, kynurenine, and the chemokines CXCL10, CCL2, CCL3 and CCL4. Of these, IL-1β had the strongest correlation with bacterial loads in the granulomas of αPD-1 treated animals. Overall, the elevated bacterial loads in the granulomas of the αPD-1 treated animals were associated with increased correlations with inflammatory mediators.

### Role of caspase 1 activation in TB exacerbation due to PD-1 blockade

IL-18 was the most differentially expressed cytokine upregulated in the granulomas of αPD-1 treated animals. Although IL-1β concentrations were not different between groups, IL-1β had the strongest positive correlation with bacterial loads in αPD-1 treated animals out of all of the molecules measured. Both of these cytokines are produced as inactive precursors that are processed into bioactive cytokines by caspase 1 inflammasomes, so we next examined caspase 1 activation in granulomas. By western blot we found that procaspase 1 was detectable in all granuloma homogenates analyzed but cleaved caspase 1 was only present in some granulomas (example blot in Figure 5f and all blots shown in Supplemental Figure 3). The p10 subunit of caspase 1, a marker of caspase activation, was detected in 11 of 23 granulomas from isotype control treated animals and 14 of 23 granulomas from αPD-1 treated animals, indicating that the presence of activated inflammasomes was not increased by PD-1 blockade (Figure 5f and Supplemental Figure 3). Thus, we did not find evidence of increased caspase 1 inflammasome activation in granulomas after PD-1 blockade.

We next asked if the presence of activated caspase 1 in granulomas was associated with higher or lower bacterial loads. In isotype control treated animals, granulomas with detectable p10 had similar bacterial loads compared to granulomas with no detectable p10 subunit (Figure 5g). In contrast, in αPD-1 treated animals, granulomas with active caspase 1 had significantly elevated bacterial loads. IL-1β levels trended higher in granulomas containing activated caspase 1 in isotype control treated animals but were significantly higher in p10+ compared to p10-granulomas from αPD-1 treated animals (Supplemental Figure 4). In contrast, there was no significant difference in IL-18 levels in p10+ versus p10-granulomas (Supplemental Figure 4). Thus, upon PD-1 blockade, caspase 1 activation is not increased, but is associated with higher levels of IL-1β and exacerbation of the infection.

Prompted by the above observations, we next sought to test the possible contribution of inflammatory caspase activation to the exacerbation of Mtb infection during PD-1 blockade using the murine model. As previously shown, PD-1 KO mice succumb rapidly to Mtb infection (Figure 5h). However, PD-1/caspase1/caspase11 triple deficient mice lived significantly longer compared to PD-1 KO mice, indicating that caspase1/11 is directly involved in the early death of PD-1 KO mice. Collectively, these data support previous reports that IFNγ and TNF contribute to increased Mtb bacterial loads during PD-1 blockade and reveal the contribution of caspase 1 driven responses as well.

### T cell motility in granulomas

PD-1 has been shown to regulate CD4 T cell motility by regulating TCR induced stop signals upon interaction with antigen presenting cells (*24, 25*), so we next examined the impact of PD-1 blockade on the positioning and intralesional T cell trafficking. First, we examined the partitioning of CD4 T cells into the infected macrophage-rich core (marked by dense CD11b staining) and the peripheral lymphocyte cuff of the granulomas. We found no difference between the percentages of CD4 T cells localizing to the infected cores of granulomas from isotype control versus αPD-1 treated macaques (Figure 6a), suggesting that PD-1/PD-L1 interactions may not be a major factor limiting the penetration of CD4 T cells into the region of the granuloma containing the infected macrophages. Although we used mCherry labeled bacteria in these experiments, the reporter expression plasmid was lost by the time of necropsy and the fluorescence could not be visualized.

**Figure 6.**
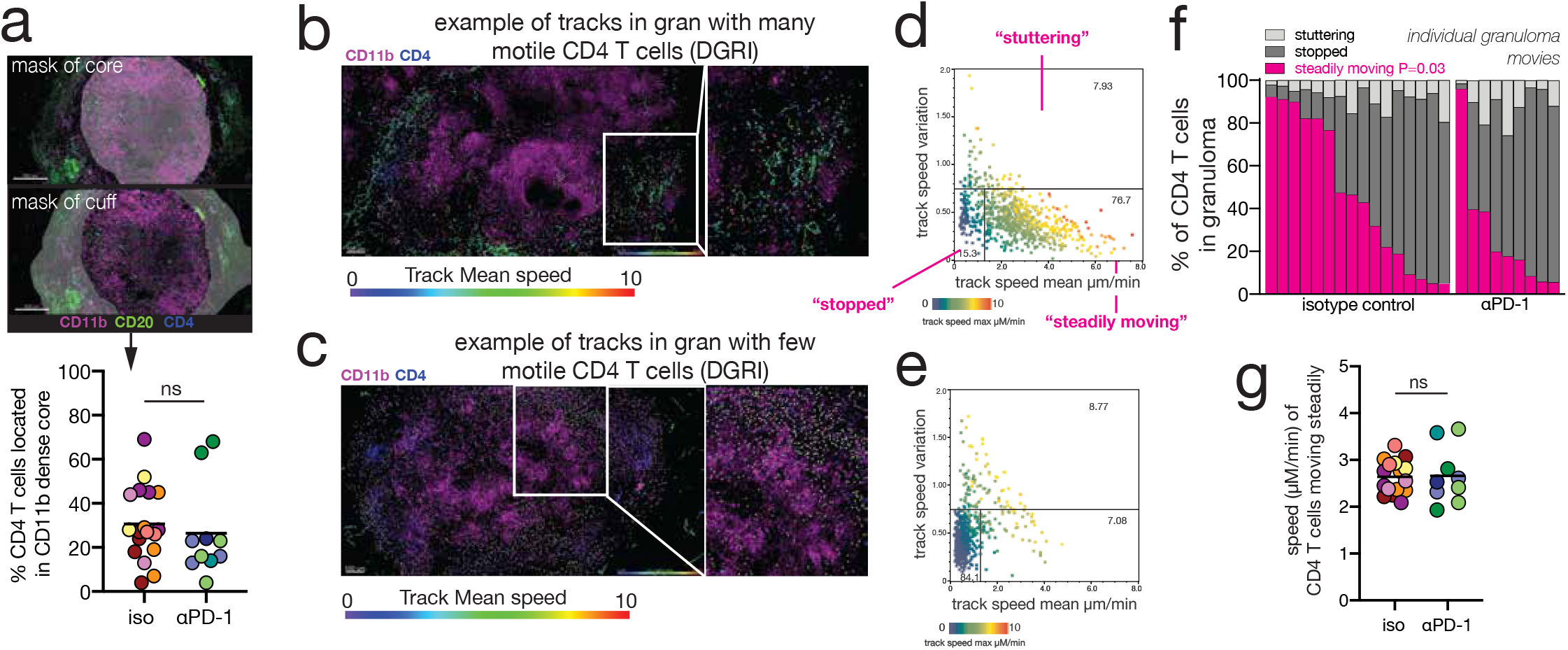
PD-1 blockade reduces CD4 T cell trafficking in granulomas. a) Example image of masking to delineate granuloma core and cuff and summary graph of percent of CD4 T cells that localized in the granuloma core. b-c) Granuloma explants were stained and imaged live to monitor T cell trafficking. Example images show CD4 T cell tracks from granulomas with (b) many motile and (c) few motile cells superimposed on the overall granuloma structure. Both example granulomas are from animal DGRI (isotype control). d-e) Plotting the mean speed and track speed variation of CD4 T cell tracks shown in the granulomas to the left identifies three basic types of movement: 1) low mean and variation in speed tracks (stopped), 2) tracks with high mean speed and low speed variation are designated (steadily moving) and 3) tracks with intermediate mean speed and high variation were designated (stuttering). Panel d corresponds to the image of the highly motile granuloma in (b) and panel (3) corresponds to the granuloma in (c) with very few motile CD4 T cells. f) Percent of each CD4 T cell movement type in each granuloma movie. Each bar represents an individual granuloma from either isotype control or anti-PD-1 treated animals. Pink bar indicates the percent of CD4 T cells that were found to be steadily moving during the movie capture. g) Average speed (micrometers/minute) of CD4 T cells. Each dot represents the average of all “steadily moving” CD4 T cells in an individual granuloma movie.

We next sought to quantify the motility of CD4 T cells in macaque tuberculomas. To do so, we performed live imaging of thick section explants of freshly isolated granulomas (Supplemental Figure 5a). To identify high quality tracks, we excluded tracks comprised of less than five individual spots, tracks that were on the top or bottom of the Z-stack, tracks whose displacement was limited by the top or bottom of the slice, and tracks that displayed significant costaining for non-T cell markers such as CD20 or CD11b (Supplemental Figure 5b). We found a high degree of heterogeneity between granulomas in the movement of CD4 T cells. Some granulomas contained many moving CD4 T cells while others displayed very few, and large differences were seen even in the same macaque (Figure 6b-c). We characterized three basic types of T cell motility in granulomas: tracks with low mean velocity and low variation in speed (stopped cells), tracks with high mean speed and low variation in speed (steadily moving cells), and tracks with intermediate mean velocity and high variability (stuttering cells) (Figure 6d-e). We specifically focused on the steadily moving CD4 T cells. Collectively, there was a significant, albeit small, trend for there to be fewer granulomas with an abundance of steadily moving cells in the αPD-1 treated animals (Figure 6f). The mean velocity of cells that were steadily moving, however, was not different in granulomas from isotype control and αPD-1 treated macaques (Figure 6g). Therefore, PD-1 blockade did not affect the speed of cells that were moving, but it resulted in a reduction in the frequency of granulomas that had abundant motile CD4 T cells.

### Association between TB exacerbation after PD-1 blockade and the gut microbiota

The composition of the intestinal microbiota has been shown to influence responsiveness to PD-1 blockade during cancer immunotherapy, and in prior work we noted an correlation between the microbiota composition and disease outcome in Mtb infected macaques (*26–30*). In the present study, 2 animals treated with αPD-1 were “non-responders” in that they did not develop increased bacterial loads or disease relative to controls, so we next asked if exacerbation of TB after PD-1 blockade was associated with the microbiota of the individual macaques. Neither Mtb infection nor PD-1 blockade resulted in changes to the diversity of the intestinal microbiota as measured by the Shannon index (Figure 7a). However, a trend was observed in which animals with lower granuloma Mtb loads clustered separately from the other animals in their treatment group (DG5F in the isotype control group and DG3X and DG2T in the αPD-1 treated group) (Figure 7b). More importantly, the pre-infection microbiotas of DG3X and DG2T were similar to each other and significantly different from the “responder” animals (Figure 7c). Among the αPD-1 treated animals, several bacterial families were specifically enriched in the responder and nonresponder animals (Figure 7d and Supplemental Figure 6). Of note, the abundance of *Ruminococcaceae* has previously been reported to associate with tumor reductions after PD-1 blockade (*29*), and here this family was strongly enriched in the monkeys that developed increased Mtb burdens after PD-1 blockade. Therefore, similar to its influence on anti-tumor responses after PD-1 blockade during cancer immunotherapy, the intestinal microbiota may also have a role in determining the outcome of PD-1 blockade during Mtb infection.

**Figure 7.**
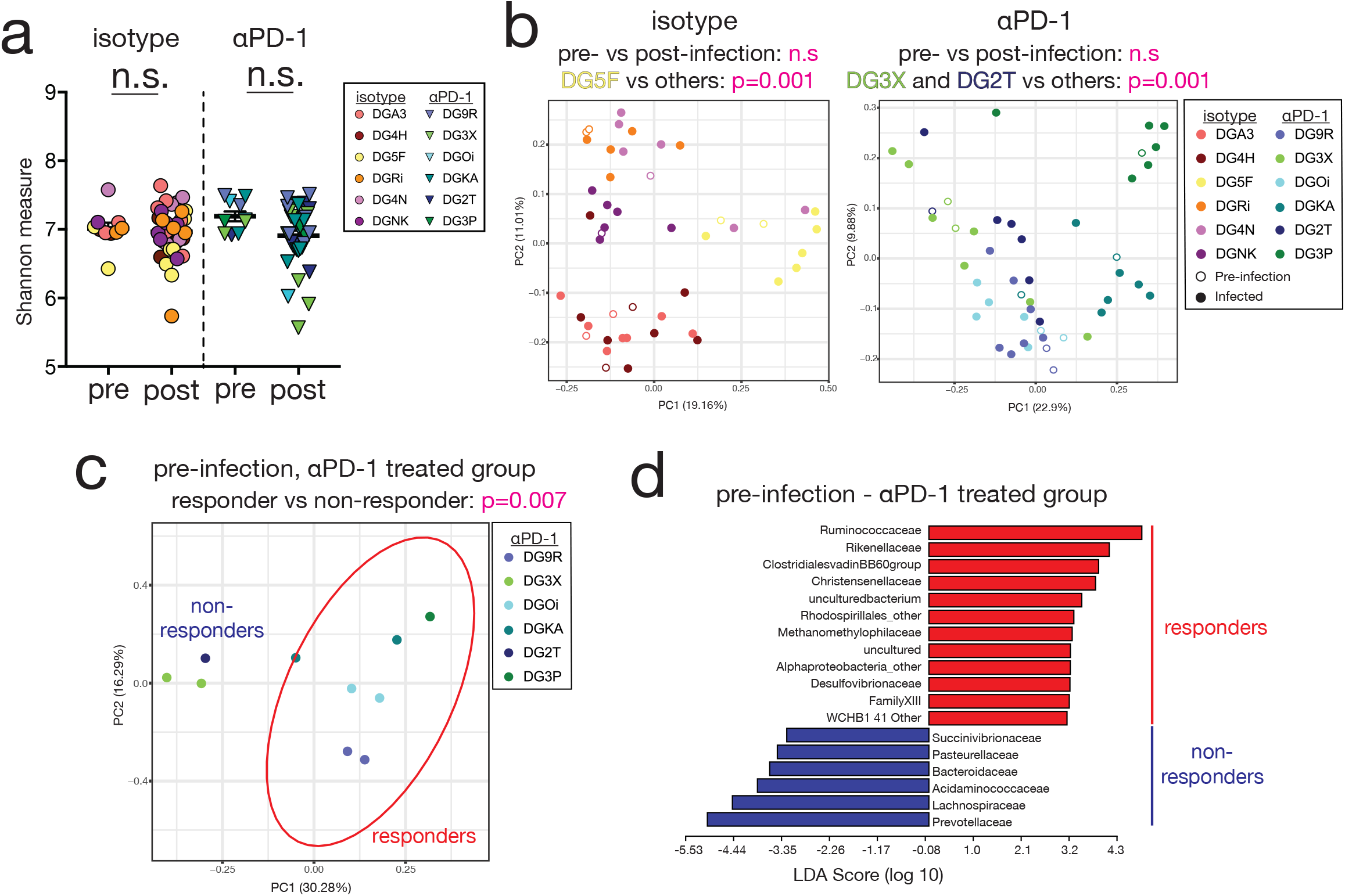
Exacerbation of Mtb infection by PD-1 blockade is associated with the composition of the pre-infection intestinal microbiota of the individual macaque. a) Alpha diversity for each sample was estimated using the Shannon index. Statistical significance between the pre- and post-infection timepoints within the isotype or αPD-1 group was calculated using Mann-Whitney U test. b) Beta-diversity analyses were performed using the Bray-Curtis dissimilarity matrix for each group and represented on a principal component (PC) plot. Each circle denotes a time point and is colored by monkey as indicated in the key with open and closed circles representing pre- and post-infection timepoints respectively. Statistical significance was calculated using PERMANOVA with 999 permutations between pre- and postinfection timepoints and animals as indicated for both study groups. DG5F in the isotype group and DG3X and DG2T in the αPD-1 group were monkeys with mild TB disease within that group and were compared to animals with severe TB in the same group. c) Beta-diversity analysis of just the pre-infection sample sequence data from the αPD1 group is represented on a PC plot. Statistical significance was calculated between animals that progressed to mild (non-responder) versus severe (responder) TB disease as previously specified. d) LEfse was used to identify differentially abundant bacterial families in the pre-infection microbiota of the non-responder versus responder monkeys in the αPD-1 group.

## DISCUSSION

These data indicate that negative regulation of immune responses is a critical aspect of host resistance to Mtb infection. The mechanisms underlying the increased Mtb bacterial loads after PD-1 blockade in infected macaques are not clear. However, IFNγ and TNF have both been previously implicated in increased growth of Mtb after PD-1 blockade (*11, 12*), and here IFNγ and TNF and IL-18 were the most dysregulated cytokines in granulomas and each positively correlated with bacterial loads after PD-1 blockade. In vitro studies have shown that stimulating Mtb infected macrophages with high concentrations of IFNγ leads to the death of the macrophage instead of the expected killing of the bacteria (*31, 32*), indicating Th1 cell function likely needs to be appropriately balanced for optimal control of the infection. We have now shown that caspase 1 activation is also required for exacerbation of Mtb infection after PD-1 blockade. Interestingly, we did not find evidence that PD-1 blockade increased caspase 1 activation in infected macaques, but only granulomas with activated caspase 1 displayed increased bacterial loads after PD-1 blockade. This raises the interesting possibility that PD-1 blockade resulted in the preferential exacerbation of Mtb infection only in the subset of granulomas with ongoing caspase 1 activation. It is not clear if the detrimental effects of caspase 1 during PD-1 blockade is due to its role in generating inflammatory cytokines or mediating pyroptotic cell death. It is also not clear why the cleaved p10 subunit of caspase 1 was only detected in a subset of granulomas in the first place. However, our findings clearly demonstrate that inflammatory pathways, normally important for host defense, are required for the exacerbation of Mtb infection after PD-1 blockade.

In mice, we have previously shown that both CD4 and CD8 T cells contributed to the early death of Mtb infected PD-1 KO mice (*10*), with CD4 T cells playing a more prominent role. Although we cannot draw conclusions regarding the contributions of CD4 and CD8 T cells to the detrimental outcome of PD-1 blockade in the present macaque study, there was a striking difference in the effect of PD-1 blockade on these two T cell subsets. The expansion and function of Mtb-specific CD8 T cells in granulomas was significantly enhanced after PD-1 blockade, while Mtb-specific CD4 T cells displayed no apparent increase in number and function. Instead, Ag-specific CD4 T cell accumulation in the airways was delayed, they upregulated high levels of CTLA-4, and displayed reduced motility in granulomas. These observations suggest that disrupting PD-1-dependent regulation may have negative consequences on some CD4 T cell functions. PD-1 primarily regulates signals through the TCR and CD28 (*33–35*), which are arguably the principle molecules T cells use to sense their environment and decide how to respond. Therefore, alterations in normal PD-1-mediated modulation of signals 1+2 should be expected to affect many aspects of T cell behavior in vivo, perhaps in unexpected ways. Counter-intuitively, it is possible that co-inhibition may be required for optimal protection against tuberculosis not only by preventing the detrimental over production of inflammatory products such as IFNγ and TNF, but also because PD-1-mediated tuning of TCR and CD28 signaling is important in allowing other normally beneficial T cell functions during chronic infection. These two possibilities are not mutually exclusive.

The timing of blockade relative to infection as well as the particular pathogen present are likely key factors in determining if PD-1 blockade will be beneficial versus detrimental in any given infectious setting. For example, during chronic LCMV infection in mice, PD-1 blockade several months after infection results in enhanced CD8 T cell responses and reductions in viral loads, while blockade in the first week of infection results in the rapid death of the animal due to CD8 T cell-mediated destruction of vascular endothelium (*36*). Here we started PD-1 blockade at the approximate time of T cell priming after Mtb infection, and it is not clear if PD-1 blockade will exacerbate tuberculosis when initiated at late stages of infection. PD-1 blockade is unlikely to exacerbate all mycobacterial infections, as among different mycobacterial species the impact of PD-1 deficiency on infection outcome varies. While PD-1 deficient mice succumb after Mtb infection, they clear BCG infection more rapidly than WT controls (*9–11, 37*). While the data indicate that PD-1 blockade is not likely to be a tenable HDT for TB, this is not the case for all infections. For example, PD-1 blockade has been show to enhance resistance to several viral (*38–40*), bacterial (*41*), parasitic (*42*) and fungal (*43, 44*) infections in mice, SIV in macaques (*45, 46*), and JC virus in progressive multifocal leukoencephalopathy in humans (*47*).

These data highlight the potential challenge of enhancing inflammatory responses as an HDT strategy for TB. The possibility of disease exacerbation seems a major concern for approaches meant to boost host responses during Mtb infection. However, there may be safe ways to stimulate protective host responses against Mtb infection by modulating other immune checkpoint molecules. For example, mice deficient in another co-inhibitory receptor, TIM-3, are more resistant to Mtb infection (*48*). A better understanding of the role of different checkpoint molecules in the regulation of host responses in tuberculosis will aid in the development of HDTs for TB.

With our findings in in macaques confirming the murine model data and accumulating clinical case reports, it seems prudent that TB should be considered a potential serious adverse event of PD-1 targeting therapies. The relative risk of tuberculosis after PD-1 blockade is not clear, but in this study four of six macaques developed increased bacterial loads after PD-1 blockade (even during infection with a less virulent strain) indicating that checkpoint blockade-associated exacerbation of tuberculosis might not be rare in Mtb infected individuals. As has been demonstrated during PD-1 targeting cancer therapies, the outcome of Mtb infection during PD-1 blockade also may be influenced by the composition of the intestinal microbiome. Based on the combined clinical and experimental data, it seems warranted that patients should be tested for Mtb exposure prior to initiation of PD-1 targeting immunotherapy and then monitored for development of active TB during treatment. Indeed, it is also increasingly clear that tuberculosis is an important concern in the development of novel immune checkpoint-targeting cancer immunotherapies, as is already the case for immunosuppressive drugs like TNF and JAK inhibitors, and as additional co-inhibitory receptors are developed as targets for cancer immunotherapy it will be important to address the possibility of TB as an adverse event.

## MATERIALS AND METHODS

### Rhesus macaques

Twelve healthy, male, 2-year-old rhesus macaques were received from the NIAID breeding colony on Morgan Island and were tuberculin skin test negative. Animals were housed in nonhuman primate biocontainment racks and maintained in accordance with the Animal Welfare Act, the Guide for the Care and Use of Laboratory Animals and all applicable regulations, standards, and policies in a fully AAALAC International accredited Animal Biosafety Level (ABSL) 3 vivarium. All procedures were performed utilizing appropriate anesthetics as listed in the NIAID DIR Animal Care and Use Committee approved animal study proposal LPD25E. Euthanasia methods were consistent with the AVMA Guidelines on Euthanasia and endpoint criteria listed in the NIAID DIR Animal Care and Use Committee approved animal study proposal LPD 25E.

### Infections and in vivo antibody blockade

Animals were infected with 30-50 colony forming units (CFU) of an mCherry-expressing *H37Rv* strain of Mtb. For infection, animals were anesthetized and 2ml of saline containing the bacteria were bronchoscopically-instilled into the right lower lung lobe. Infection dose was confirmed by plating of aliquots onto 7H11 agar plates. At weeks 2, 4, 6, 8, 10, and 13 post infection, animals were intravenously infused with 10mg/kg of body weight of either rhesus macaque IgG4 isotype control antibody (DSPR4) obtained from the NHP Reagent Resource or anti-PD-1 (humanized clone EH12 kappa variable domains with rhesus macaque kappa and IgG4 constant regions) (*49, 50*).

### PET/CT scanning procedures

Rhesus were imaged prior to infection and every two weeks after infection beginning at 4 weeks for a maximum of 8 PET/CT scans. Animals were initially anaesthetized using Glycopyrrolate (0.01 mg/kg IM) and a cocktail of Ketamine and dexmedetomidine (10mg/kg and 0.03mg/kg respectively IM) followed by insertion of an endotracheal tube. The anesthesia plane was maintained with Isoflurane (1-5% in oxygen) throughout the imaging session. [^18^F]FDG doses were determined by measuring the radioactivity content in the syringe using a dose calibrator (Biodex, New York USA). Blood glucose was measured prior to injecting [^18^F]FDG intravenously (1 mCi/kg). Injected activity and weight of the animal were recorded. During a [^18^F]FDG uptake period of 60 minutes, animals were moved to the bed of a LFER 150 PT/CT scanner (Mediso Inc, Budapest, Hungary) positioned head first, prone for imaging. While anesthetized, the animal’s heart rate, respiration, temperature, peripheral capillary oxygen saturation, and end-tidal CO_2_ were monitored. During imaging, animals received Sodium chloride 0.9% at 5-10 ml/kg/hr IV and were mechanically ventilated. A 360-projection CT scan of the lungs was acquired while the animal was maintained in a breath hold for about 50 seconds. The parameters of the CT scan were: 80 kVp, 0.820 mA and 80 ms/projection. The resulting CT image was inspected for potential movement or mispositioning and repeated if necessary. The CT data was used to generate a material map for the reconstruction of the corresponding PET image. A 20-minute PET data set was acquired (Coincidence mode 1-9) with the identical field of view during mechanical ventilation. Following imaging, animals were administered Antisedan (0.3mg/kg IM) to reverse sedation and returned to their home cages. The raw CT and PET data were reconstructed (CT: Voxel size 0.5X0.5X0.5 mm Butterworth filter, PET: 3D Tera-Tomo reconstruction 8 iterations 9 subsets Voxel size 0.8 mm with random, attenuation and scatter correction) using the Nucline software (Mediso, Inc, Budapest, Hungary).

### PET and CT data analysis

PET and CT DICOM files were co-registered using MIM Maestro (v. 6.2, MIM Software Inc, Cleveland, Ohio). A lung volume of interest (VOI) was defined on the CT image using methods previously described including adjusting the VOI away from non-lung surfaces by shrinking the VOI by 0.5 mm (*51*). For CT analysis, a lower threshold of −400 HU was applied and the remaining dense VOI was inspected to confirm the inclusion of only areas of granulomatous disease. For PET analysis, the adjusted CT VOI was transferred to the PET image and a lower threshold of 2.5 SUVbw was applied to remove background FDG uptake. The remaining FDG uptake in the VOI was considered tubercular disease-related (*51, 52*). Manual removal of nonspecific [^18^F]FDG uptake was occasionally required in proximity of large blood vessels and the diaphragm. The software generated tables with all the values recorded for the selected PET and CT volumes for the serial scans. Two readers blind to treatment assignment independently performed image analysis above for each animal. If there was more than 5% disagreement, a third read was included and the disease findings from each assessment were averaged. Threedimensional projections were generated using Osirix v 5.9 software (Pixmeo, Geneva, Switzerland).

### Cell isolations and stimulations

Blood samples were collected in EDTA tubes and PBMCs were isolated by Ficoll-Paque (GE Life Sciences) density centrifugation. Broncho-alveolar lavage (BAL) samples were passed through a 100 μm cell strainer, pelleted, and counted for analysis. Lymph nodes and spleens were dissociated using a GentleMACS Tissue Dissociator (Miltenyi Biotech). Granulomas were individually resected from the lungs and samples used for flow cytometry analysis were pushed through a 100 μm cell strainer. Aliquots from all samples were serially diluted and plated on 7H11 agar plates for CFU quantification. Cells were stimulated for 6 hours at 37 °C in X-vivo 15 media supplemented with 10% FCS with MHC-I and MHC-II (MTB300) Mtb peptide megapools (1 μg/ml and 2 μg/ml respectively) in the presence of brefeldin A and monensin (eBioscience).

### Flow cytometry

Fluorochrome-labeled antibodies used for flow cytometric analysis are listed in Table S1. Surface antigens and dead cells were stained in PBS + 1% FCS + 0.1% sodium azide for 30 minutes at 4 °C. For intracellular cytokine and transcription factor staining, cells were fixed and permeabilized with the Foxp3 Transcription Factor Staining Buffer Kit (eBioscience) and stained for 30 minutes at 4 °C. Samples were acquired on a FACSymphony (BD Biosciences), and data were analyzed using FlowJo 10 (Treestar). Lymphocytes were gated on by the exclusion of dead cells and doublets and were CD3^+^.

### Multiplex cytokine analysis

Granuloma homogenates and plasma samples were sterile filtered and analyzed for protein concentrations using the Invitrogen Cytokine and Chemokine 30-Plex ProcartaPlex kit (Thermo Fisher). Samples were acquired on a Magpix with xPonent software (Luminex Corporation). A kynurenine ELISA was also performed using assay instructions on the homogenate samples (Labor Diagnostika Nord). Statistical analysis of this data is described below.

### Western Immunoblotting

Total protein from granuloma supernatants were normalized by dilution in 1x Tris-buffered saline (TBS). THP-1 monocytes were plated in Opti-MEM (Gibco) + 1% FCS and treated with 500 ng/mL Pam_3_CSK_4_ (InvivoGen) for 4 hours and 10 μM nigericin (InvivoGen) for an additional 1 hour. 1.2×10^6^ cells were lysed in 300 μL RIPA buffer supplemented with cOmplete Protease Inhibitor Cocktail (Roche) on ice for 30 minutes and passed through 30 μm polyethylene filter spin columns (Pierce) to shred genomic DNA. 2mL cell-free THP-1 supernatant was concentrated to 100 μL by methanol-chloroform precipitation (*53*). All samples were denatured by addition of SDS sample buffer (final concentration 70 mM SDS) and boiling for 5 minutes. 8 ng of granuloma protein was loaded per lane into Any kD Mini-PROTEAN TGX gels (Bio-Rad), separated by SDS-PAGE and transferred to 0.2 μm nitrocellulose membranes (Bio-Rad). Transfer and loading were validated by staining membranes in 0.2% Ponceau S solution (Sigma-Aldrich) for 2 minutes. Membranes were blocked in 5% skim milk in TBS + 0.1% Tween 20 (TBS-T) for 1 hour at room temperature before incubation in primary antibody overnight at 4°C (rabbit-α-caspase-1, Abcam #ab179515, 1:1000). Membranes were washed in TBS-T, incubated in secondary antibody for 1 hour at room temperature (donkey-α-rabbit-HRP, Jackson ImmunoResearch #711-035-152, 1:10,000; mouse-α-actin-HRP, Santa Cruz #sc-47778, 1:10,000), washed and coated in Immobilon Western Chemiluminescent HRP Substrate (Millipore). Immunoreactivity was imaged using either Amersham Hyperfilm ECL (GE Healthcare) and an X-OMAT 2000A processor (Kodak) (caspase-1) or a ChemiDoc Touch Imaging System (BioRad) (actin).

### Confocal imaging and analysis

Granulomas were embedded in RPMI containing 2% agarose and sliced into 300–350 μm sections using Leica VT1000 S Vibrating Blade Microtome (Leica Microsystems, Exton, PA). Tissue sections were stained with fluorescently labeled antibodies specific to CD4, −CD8, −CD11b, −CD20 or PD-1 of choice (eBioscience) for 2 h on ice. After staining sections were washed 3 times and cultured in complete lymphocyte medium (Phenol Red-free RPMI supplemented with 10 % FBS, 10 % rhesus plasma, 25 mM HEPES, 2 mM L-glutamine, 1% Sodium Pyruvate, and 1% penicillin and streptomycin) in humidified incubator at 37° C. Sections were imaged using Leica SP5 inverted 5 channel confocal microscope equipped with an Environmental Chamber (NIH Division of Scientific Equipment and Instrumentation Services) to maintain 37 °C and 5% CO_2_. Microscope configuration was set up for four-dimensional analysis (*x, y, z, t*) of cell segregation and migration within tissue sections. Diode laser for 405 nm excitation; Argon laser for 488 excitation, DPSS laser for 561; and HeNe lasers for 594 and 633 nm excitation wavelengths were tuned to minimal power (between 1 and 5 %). Mosaic images of complete granuloma sections were generated by acquiring each *xy* plane with 10 – 50 μm *z* spacing every 10 seconds for 3 – 4 hours using motorized stage to cover the whole section area and assembled into a tiled image using LAS X software (Leica Microsystems, Exton, PA). Motion artefacts, 3D alignment and thermal drift of sequential z-sections was corrected using cross correlation algorithm in Huygens Pro software package (version 20.04.0p2, Scientific Volume Imaging BV, Hilversum, The Netherlands). Post-acquisition mages were processed using Imaris software (version 9.5.1, Bitplane AG, Zurich, Switzerland) for quantification and visualization. Localizations of the cells in granulomas were identified based on fluorescence intensity using spot function of Imaris and divided into different regions based on localization using surface 3D reconstruction and channel masking method. Multiparameter track data generated by the Imaris analysis of granuloma movies were imported in FlowJo to facilitate visualization and exploration of the data. Data gated in FlowJo were re-imported into Imaris using custom generated python script for spatial analysis to further localize and visualize cell subpopulations.

### Microbiota analyses

Fecal samples were collected at the timepoints indicated in Figure 1a and frozen at −80 ° C. Only one pre-infection sample was collected for monkeys DG2T, DG3P, DG4N and DGNK and the week 13 post-infection time point was not collected from monkey DGOi. At the end of the experimental timeline, DNA was extracted from 0.04-0.05 g of stored fecal material using QIAamp Fast DNA stool Mini kit (Qiagen, Hilden, Germany) as previously described (*54*). The V4 region of the 16s RNA gene was amplified with primers 5’-TCGTCGGCAGCGTCAGATGTGTATAAGAGACAGGTGCCAGCMGCCGCGGTAA-3’and 5‘-GTCTCGTGGGCTCGGAGATGTGTATAAGAGACAGGGACTACHVGGGTWTCTAAT-3’ and sequenced on an Illumina MiSeq Platform as previously described (*54*). The raw fastq data were demultiplexed and the reads were processed, denoised and filtered for chimeras using the DADA2 algorithm implemented in QIIME2 version 2-2018.4 (*55*) resulting in an average of 64,134 reads/sample. Taxonomic classification was performed using Silva database release 132 (*56*). The week 7 time point from monkey DGRi was dropped from further analysis as this sample sequenced poorly. Alpha and beta-diversity analyses on read data rarefied to a depth of 44,000 reads were performed using the Shannon and Bray-Curtis dissimilarity indices respectively. Linear discriminant analysis (LEfSe) was used to identify differentially abundant taxa between the groups compared and taxa with a linear discriminant score (LDA) of >2 and p-value < 0.05 were considered significantly different (*57*).

### Statistical analysis

Biomarkers levels were log10 transformed and normalized by mg of protein. Limit of detection values were subtracted. Data were compared between the study groups using the Mann-Whitney *U* test (two-group comparisons) to identify the biomarkers that were statistically significant between groups. P-values were adjusted for multiple measurements using Holm-Bonferroni’s method when appropriate. Hierarchical cluster analyses (Ward’s method), with 100X bootstrap of z-score normalized data were employed to depict the overall expression profile of indicated biomarkers in the study groups.

Sparse canonical correlation analysis (sCCA) modelling was employed to assess whether combinations of soluble mediators could discriminate between isotype control and anti-PD1 groups (R script: https://www.jstatsoft.org/article/view/v023i12). The sCCA algorithm was chosen because many variables were studied, and it allows for us to test whether experimental groups could be distinguished based on the overall expression profile of all the markers measured as well as mycobacterial loads (*M. tuberculosis* CFU values per mg of protein). This approach reduces dimensionality for two co-dependent data sets (biomarker profile and PD-1 blockade) simultaneously so that the discrimination of the clinical endpoints represents a combination of variables that are maximally correlated. Thus, trends of correlations between parameters in di erent clinical groups rather than their respective distribution within each group are the key components driving the discrimination outcome. In our CCA model, simplified and adapted from previously reported investigations of biomarkers for TB diagnosis (*58, 59*), linear regression graphs represent coefficients from different combinations of plasma factors and baseline characteristics. In the biomarker profile dataset, we included values of all the inflammatory marker variables. Correlations were examined using the Spearman rank test. In the network analysis based on Spearman correlations, nodes represent each given marker and lines represent statistically significant correlations (correlation coefficient [rho] > ±.5 and P < .05).The statistical analyses were performed using JMP 14 (SAS, Cary, NC) and Prism 7.0 (GraphPad Software, San Diego, CA) and R statistical software.

## Acknowledgments

We are grateful to Dr. Rama Rao Amara for advice regarding dosing regimen of αPD-1. We thank Nannan Zhu, David Stephany and all staff members of the National Institutes of Allergy and Infectious Diseases, Comparative Medicine Branch (CMB) Animal Biosafety Level 3 facility for their technical support. This work was supported by the IRP of the NIAID. GJF is supported by P01AI056299 and R37AI112787. DLB has patents on the PD-1/PD-1 pathway. GJF has patents/pending royalties on the PD-1/PD-L1 pathway from Roche, Merck MSD, Bristol-Myers-Squibb, Merck KGA, Boehringer-Ingelheim, AstraZeneca, Dako, Leica, Mayo Clinic, and Novartis. GJF has served on advisory boards for Roche, Bristol-Myers-Squibb, Xios, Origimed, Triursus, iTeos, NextPoint, IgM, Jubilant and GV20. GJF has equity in Nextpoint, Triursus, Xios, iTeos, IgM, and GV20. Ziv Yaniv was supported by BCBB Support Services Contract HHSN316201300006W/HHSN27200002.

---

**Supplemental Figure 1. Impact of PD-1 blockade on T_regs_, MAIT cells and γδ T cell frequencies in Mtb infected macaques.**

a-c) Frequencies of foxp3+ CD4 T cells in the a) blood, b) BAL, and granulomas. d-f) Frequencies of CD3+ cells that are Vα7.2+CD161+CD8+ MAIT cells in the d) blood, e) BAL and f) granulomas. g-i) Frequency of CD3+ cells that are γδ T cells in the g) blood, h) BAL, and i) granulomas.

**Supplemental Figure 2. PD-1 blockade increases plasma levels of CD40L and CCL3.**

As done in Figure 5, soluble mediators were also measured in plasma. Only a) CD40L and b) CCL3 were both detectable and different between groups. c) All measurements from all time points were pooled for a correlation analysis between soluble CD40L and CCL3.

**Supplemental Figure 3. Detection of activated caspase 1 in granuloma homogenates.** Cleaved caspase-1 p10 was measured by Western blot analysis of granuloma homogenates. Data are representative of all samples analyzed.

**Supplemental Figure 4. Association of IL-1β and IL-18 levels with caspase 1 activation in granulomas.**

a) IL-1β and b) IL-18 concentrations in granuloma homogenates that contained a detectable activated caspase 1 p10 bands as shown in Supplemental Figure 2.

**Supplemental Figure 5. Live imaging of macaque pulmonary granulomas.**

a) Diagram of workflow for acquisition of movies. Immediately after resection granulomas were embedded in agarose, cut on a vibratome, stained with antibody cocktails, and images acquired on a confocal microscope. b) Multiparameter track data were imported into Flowjo to facilitate exclusion of artifacts and low-quality tracks. The “gating” strategy is shown. These cell populations were spatially recognized following FlowJo gating and script re-importing cell tracks back into original images in Imaris. A python script was used to select and duplicate gated tracks from FlowJo in every step of the gating process using the specific track ID. This allowed us to see and track the gated cell movement directly in the tissue and use Imaris tools to further analyze resulting tracks.

**Supplemental Figure 6. Correlation between pre-infection intestinal microbiota abundance and lung mycobacterial load at necropsy**

Mean relative abundance of certain bacterial families in the intestinal microbiota of monkeys in the αPD-1 group prior to infection are plotted are against the mean CFU/granuloma from the lungs of the same monkeys at necropsy. Spearman correlation analyses and linear regression tests were performed for each association and if significant are indicated on the top left and bottom right of each graph respectively.

**Table S1:** Flow cytometry panels. List of flow cytometry panels used in this study.

|  | **Antibody/Dye** | **Clone** | **Fluorophore** |
| --- | --- | --- | --- |
| **Panel 1** | IFNγ (in) | B27 | BUV395 |
|  | CD4 (ex) | SK3 | BUV496 |
|  | CD28 (ex) | CD28.8 | BUV737 |
|  | CD8 (ex) | SK1 | BUV805 |
|  | TNF (in) | Mab11 | BV421 |
|  | CD40L (in) | 24-31 | BV510 |
|  | IL-2 (in) | MQ1-17H12 | BV605 |
|  | CD95 (ex) | DX2 | BV711 |
|  | CD3 (ex) | SP34-2 | BV786 |
|  | PD-1 | 913429 | AF488 |
|  | CD153 (in) | 116614 | PE |
|  | IL-17A (in) | eBIO64DEC17 | Pe/Cy7 |
|  | Eomes (in) | WD1928 | APC |
|  | live/dead dye | NA | Fixable Viability Dye 780 |
| **Panel 2** | TNF (in) | Mab11 | BUV395 |
|  | CD4 (ex) | SK3 | BUV496 |
|  | CD69 (ex) | FN50 | BV711 |
|  | CD8 (ex) | SK1 | BUV805 |
|  | Granzyme B (in) | GB11 | BV421 |
|  | Ki67 (in) | B56 | BV510 |
| | IFNγ (in) | B27 | BV605 |
|  | CD127(ex) | eBIORD3 | SB702 |
|  | CD3 (ex) | SP34-2 | BV786 |
|  | Perforin (in) | Pf-344 | FITC |
|  | LIGHT (in) | T5-39 | PE |
|  | CTLA-4 (in) | 14D3 | Pe/Cy7 |
|  | PD-1 | 913429 | AF647 |
|  | live/dead dye | NA | Fixable Viability Dye 780 |
| **Panel 3** | TCRgd (ex) | B1 | BUV395 |
|  | CD4 (ex) | SK3 | BUV496 |
|  | CD8 (ex) | SK1 | BUV805 |
|  | PD-1 (ex) | EH12 | BV421 |
|  | Va7.2 (ex) | 3C10 | BV605 |
|  | CD3 (ex) | SP34-2 | BV786 |
|  | Foxp3 (in) | 150D | AF488 |
|  | PLZF (in) | MAGS2F7 | PE |
|  | CD161 (ex) | HP3G10 | Pe/Cy7 |
|  | PD-1 | 913429 | AF647 |
|  | live/dead dye | NA | Fixable Viability Dye 780 |

**Supplemental Movie 1. Example live imaging movie of thick section granuloma explant.**

Movie from DGRi. Blue = CD4, green CD20, and magenta CD11b.

