## Supplementary figures and images for "PD-1 blockade exacerbates *Mycobacterium tuberculosis* infection in rhesus macaques"

### Supplemental Figure 1

#### Supplemental Figure 1

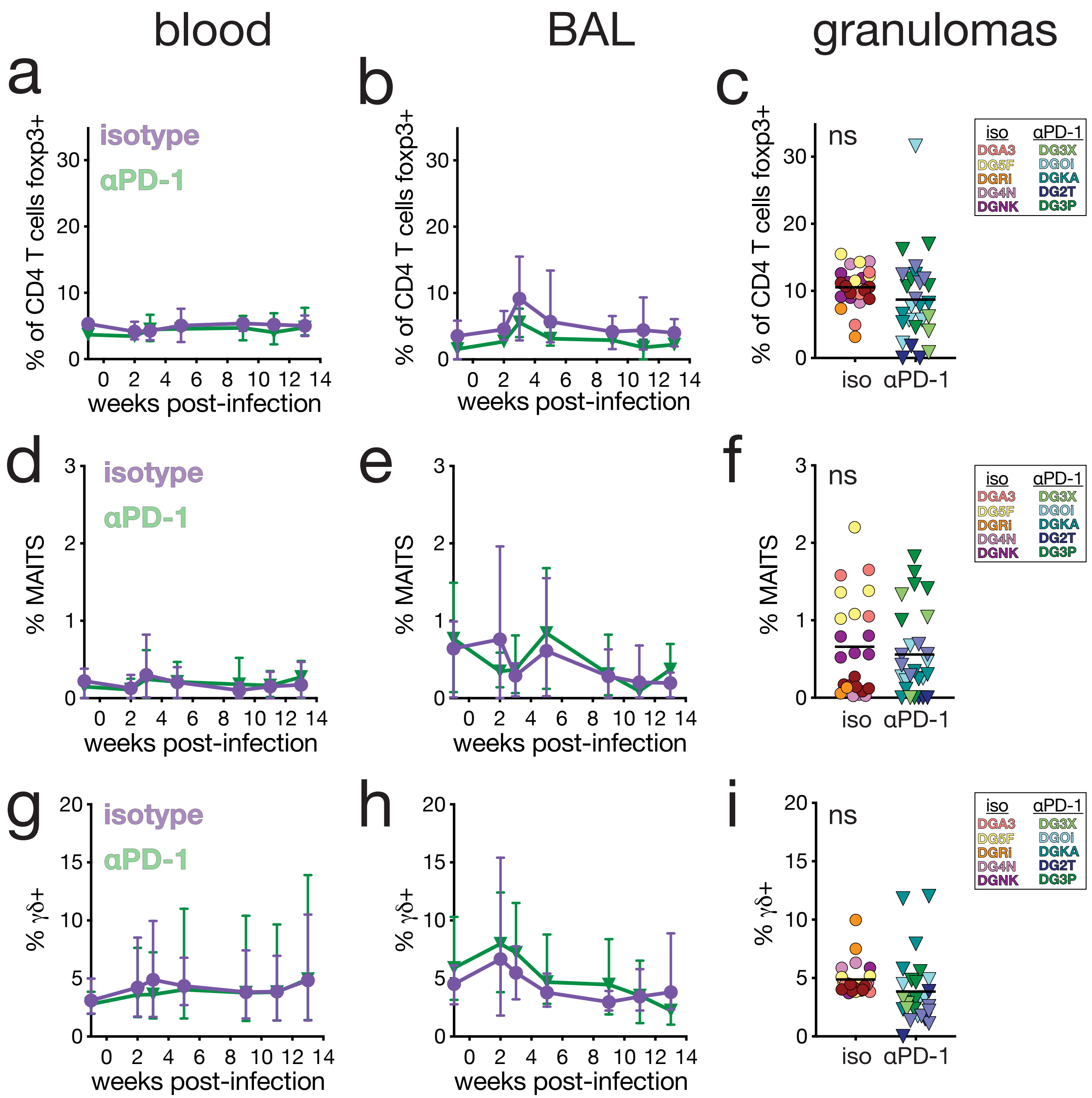

### Supplemental Figure 2

#### Supplemental Figure 2

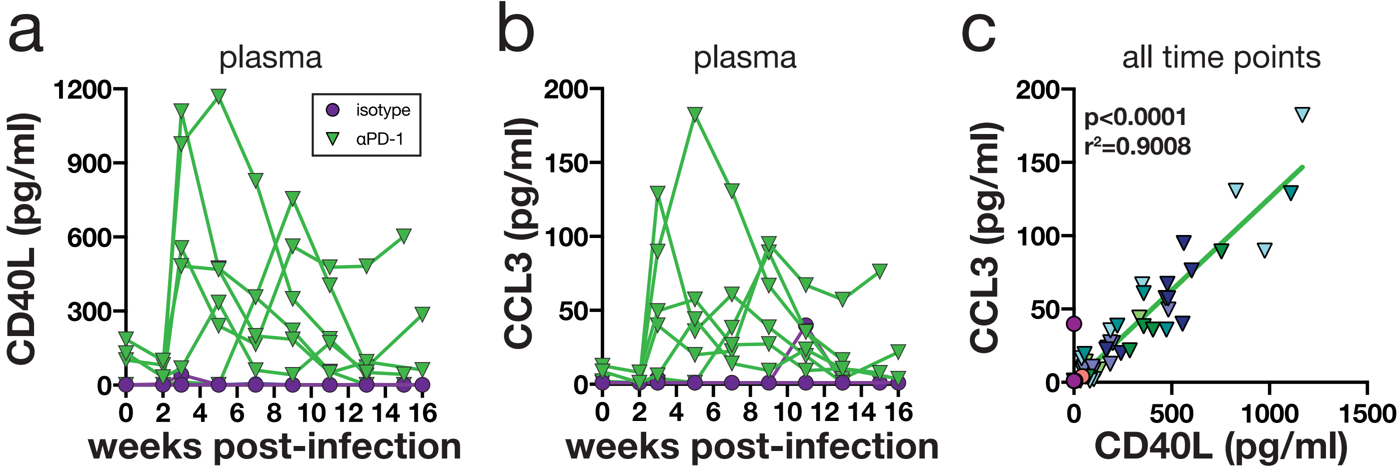

### Supplemental Figure 3

#### Supplemental Figure 3

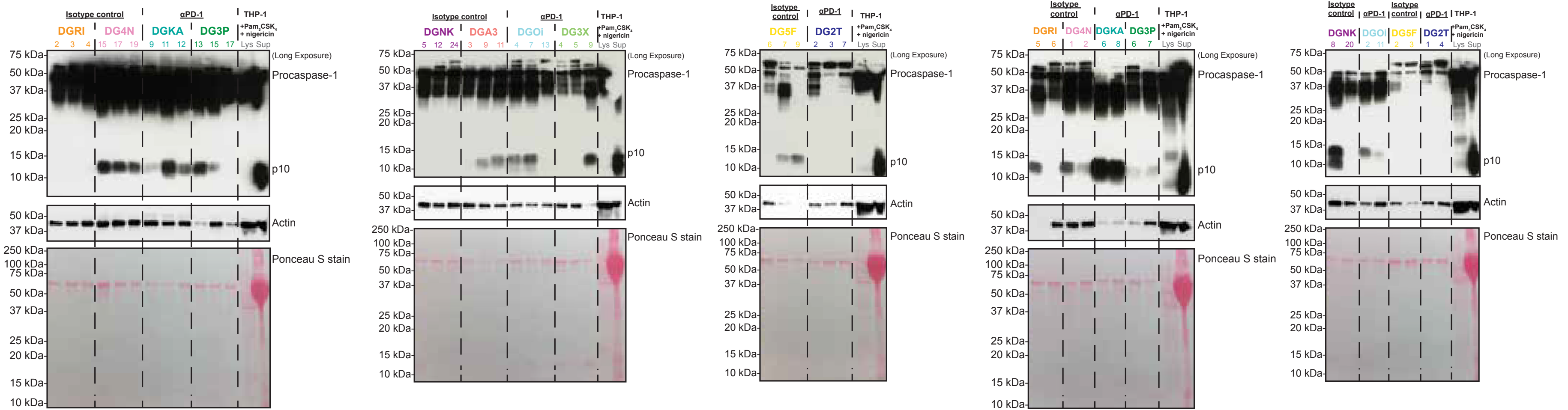

### Supplemental Figure 4

#### Supplemental Figure 4

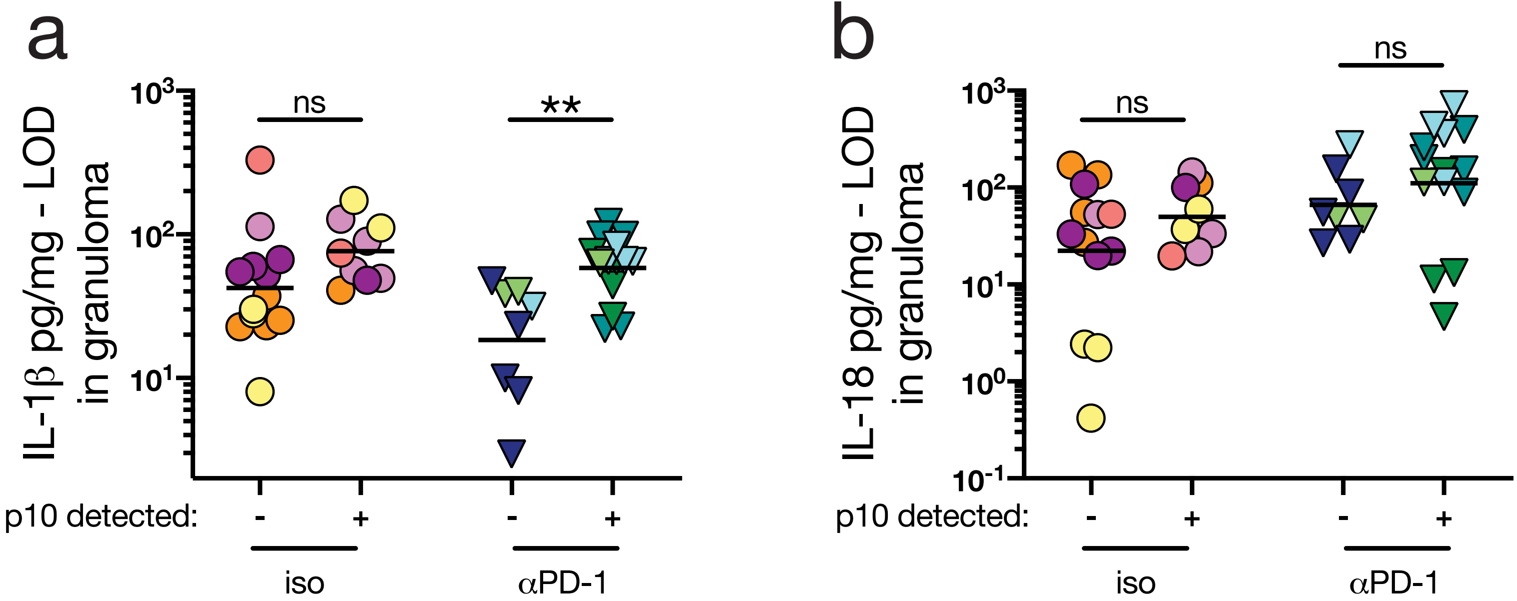

### Supplemental Figure 5

#### Supplemental Figure 5

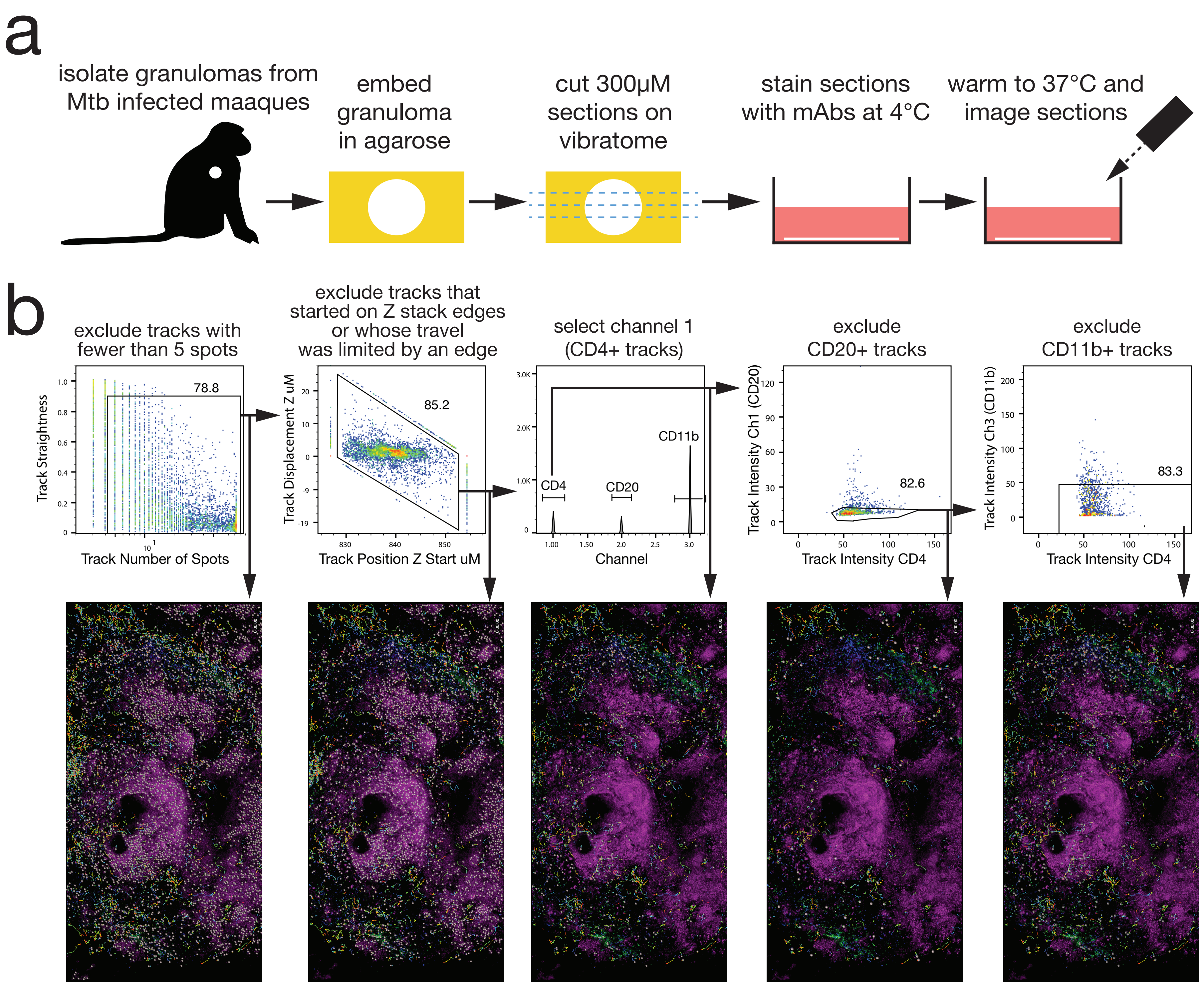
