## Supplemental Figure 6 for "PD-1 blockade exacerbates *Mycobacterium tuberculosis* infection in rhesus macaques"

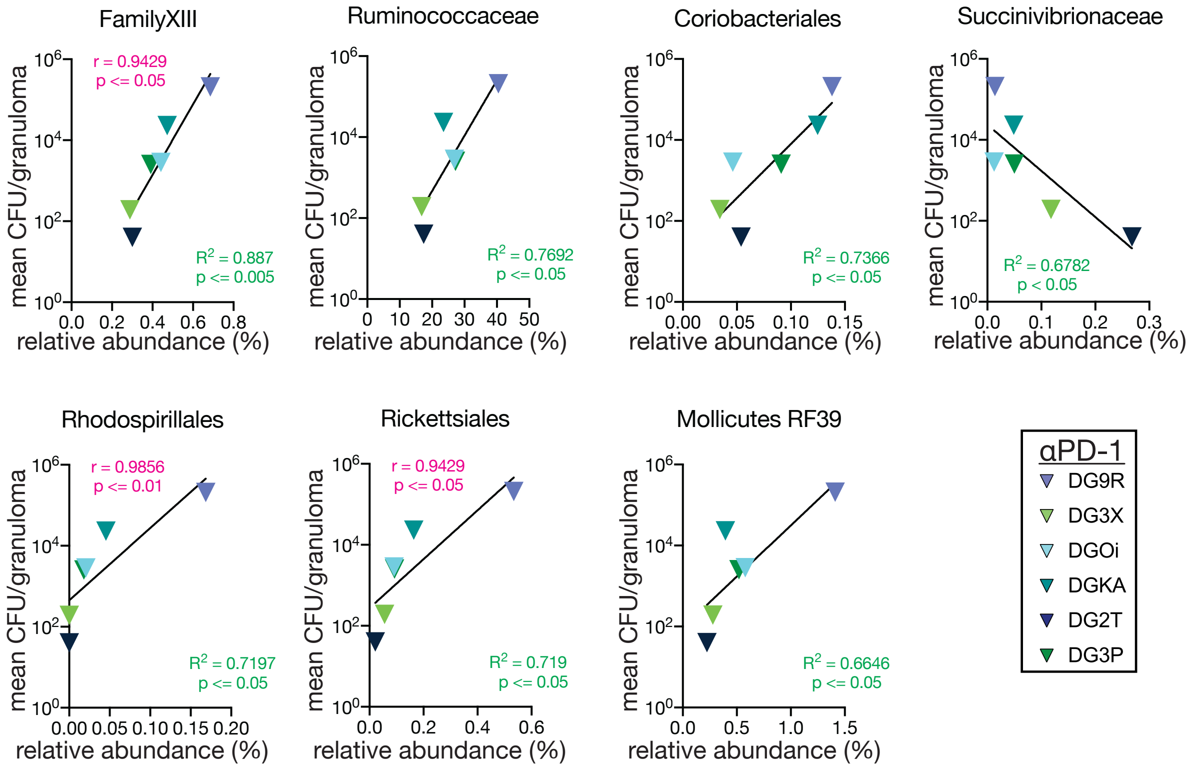


Top left: Spearman correlation stats

Bottom Right: Linear regression stats
